# Phylogeny-aware Identification and Correction of Taxonomically Mislabeled Sequences

**DOI:** 10.1101/042200

**Authors:** Alexey M. Kozlov, Jiajie Zhang, Pelin Yilmaz, Frank Oliver Glöckner, Alexandros Stamatakis

## Abstract

Molecular sequences in public databases are mostly annotated by the submitting authors without further validation. This procedure can generate erroneous taxonomic sequence labels. Mislabeled sequences are hard to identify, and they can induce downstream errors because new sequences are typically annotated using existing ones. Furthermore, taxonomic mislabelings in reference sequence databases can bias metagenetic studies which rely on the taxonomy. Despite significant efforts to improve the quality of taxonomic annotations, the curation rate is low because of the labour-intensive manual curation process.

Here, we present SATIVA, a phylogeny-aware method to automatically identify taxonomically mislabeled sequences (“mislabels”) using statistical models of evolution. We use the Evolutionary Placement Algorithm (EPA) to detect and score sequences whose taxonomic annotation is not supported by the underlying phylogenetic signal, and automatically propose a corrected taxonomic classification for those. Using simulated data, we show that our method attains high accuracy for identification (96.9% sensitivity / 91.7% precision) as well as correction (94.9% sensitivity / 89.9% precision) of mislabels. Furthermore, an analysis of four widely used microbial 16S reference databases (Greengenes, LTP, RDP and SILVA) indicates that they currently contain between 0.2% and 2.5% mislabels. Finally, we use SATIVA to perform an in-depth evaluation of alternative taxonomies for Cyanobacteria.

SATIVA is freely available at https://github.com/amkozlov/sativa.

## I. Introduction

Taxonomy is the science of classifying and naming groups of organisms, usually based on shared characteristics and/or presumed natural relatedness. Taxonomies are of fundamental importance for biological, medical and environmental research. Furthermore, they play a key role in areas such as invasive species management (1) or trade facilitation (2).

Although first attempts to classify living organisms can be traced back to antiquity (e.g., Aristotle), modern taxonomy has its origin in the work of Carl Linnaeus. His unique binomial system, that is still being used today, standardized species naming across all domains of life, from bacteria to animals. However, taxonomic classification methods have witnessed a paradigm change over the last decades, driven by progress in molecular biology and bioinformatics. Instead of exclusively relying on, e.g., morphological or physiological similarities among organisms, taxonomists now typically also take into account their phylogenetic relationships as inferred from molecular data (DNA or amino acid sequences).

While molecular phylogenies offer a more robust framework for devising taxonomies, they *do* exhibit some potential pitfalls. Firstly, a phylogeny essentially represents an evolutionary hypothesis, which is subject to the amount and quality of sequence data, the alignment quality as well as the inference method and parameters. Therefore, taxonomies that are based on phylogenies need to be updated as new sequences and methods become available. This is often not the case. Furthermore, problems inherent to molecular data, such as chimeric and/or low quality sequences (3; 4), may affect phylogenetic inferences. Finally, human error is always present; wrong cultures for organisms or mislabels in public databases can further complicate the phylogenetic analysis and the subsequent taxonomic annotation.

Microbial organisms, collectively Bacteria, Archaea, and microscopic Eukaryota, represent the most diverse group of living organisms. Unfortunately, microbial organisms are notoriously difficult to characterize as less than 1% of the microbes have been successfully cultivated so far (5). Therefore, a major breakthrough in the field of microbial taxonomies was the use of the ribosomal rRNA gene (particularly the small subunit thereof, the SSU which is called 16S rRNA for Bacteria and Archaea and 18S rRNA for Eukaryota). Carl Woese recognized that molecular evidence would revolutionize the field of bacterial phylogeny and taxonomy, since the approach could replace the rather uninformative comparative anatomy and physiology approaches (6) used at that time. Molecular methods allowed researchers to elucidate the evolutionary relationships among distant microbial lineages, leading to a unified classification of life into three domains (the ‘three-domain system’).

Norman R. Pace and colleagues (7) further extended Woeses work through the development of environmental PCR, enabling the amplification of rRNAs directly from environmental samples and assessments of microbial diversity at a molecular scale (8; 9). Moreover, recent studies correlated changes in the gut microbial composition with human conditions such as obesity, diabetes, and inflammatory bowel disease (10; 11; 12). The prerequisite for carrying out such environmental studies is the availability of a reliable taxonomic classification of the environmental sequences. In turn, this requires a stable and well-curated taxonomy for the corresponding reference database sequences.

From a computational point of view, there are two distinct approaches to microbial taxonomy. First, knowledge bases such as the List of Prokaryotic names with Standing in Nomenclature (LPSN, (13)), the Index Fungorum (www.indexfungorum.org), or the Integrated Taxonomic Information System (www.itis.gov) provide information about taxonomy and nomenclature. Second, sequence databases such as SILVA (14), RDP-II (15), Greengenes (16), EzTaxon (17), PR2 (18), or UNITE (19) maintain reference collections of taxonomically annotated sequences. Mislabels are especially problematic for sequence databases, where incorrect annotations are commonly associated with the classification algorithm being used or with low-quality sequence data. However, taxonomic errors may also occur in knowledge bases, for example due to initial misidentification of a species, or due to insufficient external sequence data for correctly arranging taxa. These errors are then propagated to the sequence databases which rely on the knowledge bases for their taxonomies. The iterative update procedures can further amplify the problem, since potentially incorrect annotations of existing sequences are used to classify new sequences. Of course, such errors can be eliminated by means of manual curation and continuous re-assessment of old classifications based on the new data. However, growing database size makes this approach less practical: some erroneous annotations might escape the curator’s attention, and thus persist in the database and propagate into further releases (see Supplementary Methods for an example). Therefore, we think that computational methods based on *de novo* phylogenies, such as `tax2tree` (16) and the method proposed here, can be of great value for preventing error propagation.

For some organism groups, a community-driven approach to curation has proved to be successful. Notably, UNITE provides a web platform for third-party annotation of fungal ITS sequences (20). Within such a system, work sharing and improved support via appropriate software allow to speed up curation substantially (21). However, this approach is subject to the willingness of the respective community to invest time and effort into taxonomic curation. Although changing taxonomic labels *per se* is fairly easy in systems like UNITE, the most time-consuming part still remains: the identification of problematic sequences as well as coming up with the new, corrected labels for them. Therefore, we believe that tools offering automatic recommendations for those two fundamental tasks will be beneficial for online and offline curation alike.

Here, we propose a novel method for identifying putative mislabels in taxonomies. Motivated by the current phylogeny-aware approach to taxonomy, we consider topological incongruence between the taxonomic and the phylogentic tree as an indication that some of the sequences might be mislabeled. Hence, we use the Evolutionary Placement Algorithm (EPA) (22) to identify sequences whose taxonomic and phylogenetic placements are inconsistent.

## II. MATERIALS AND METHODS

### A. SATIVA pipeline for taxonomy curation

We implemented our taxonomic curation method in a pipeline called SATIVA (Semi-Automatic Taxonomy Improvement and Validation Algorithm, see Fig. 1). It is opensource and freely available under https://github.com/amkozlov/sativa. In this Section, we briefly describe the method and provide some implementation details.

**Fig. 1:**
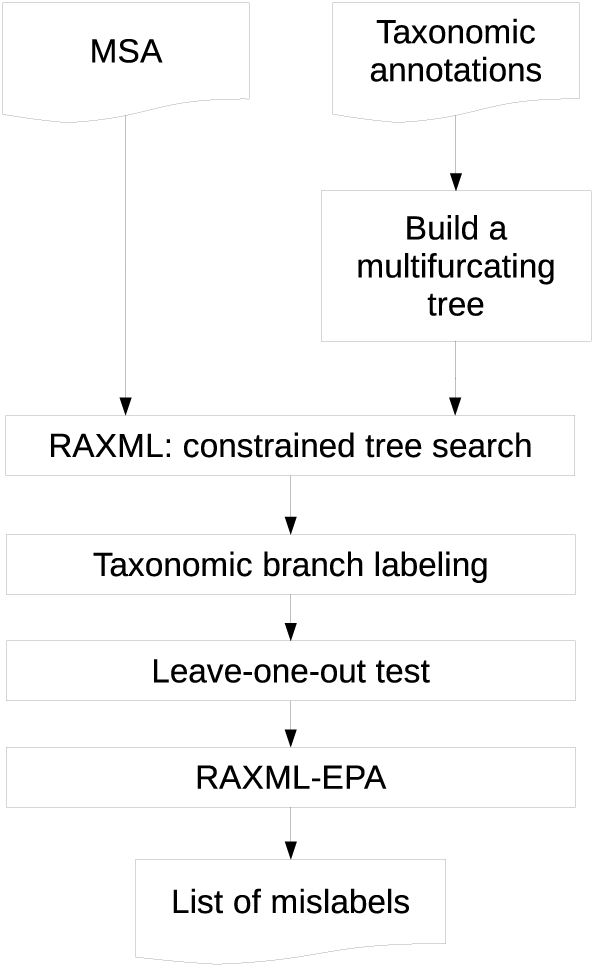
SATIVA processing workflow.

#### 1) Building a taxonomically-labeled reference tree.

Using an aligned set of sequences with taxonomic annotations, we initially build a rooted, multifurcating tree that represents the underlying taxonomy. In this tree (which we call *taxonomic tree* henceforth), leaf nodes correspond to the sequences, and inner nodes to higher taxonomic ranks such as genus and family. Then, we use RAxML (23) to perform a Maximum Likelihood (ML) tree inference using the taxonomic tree as a topological constraint. Thereby, we obtain a strictly bifurcating tree (a *reference tree*) that is fully congruent with the original taxonomic tree. Further, we label each inner node of the strictly bifurcating reference tree by the lowest common rank of its corresponding child nodes (see Fig. 2, A). For instance, given annotations (*’Escherichia’, ‘E.coli’*) and (*’Escherichia’, ‘E.albertii’*) at the child nodes, the parent node will be labeled as (*’Escherichia’*).

**Fig. 2:**
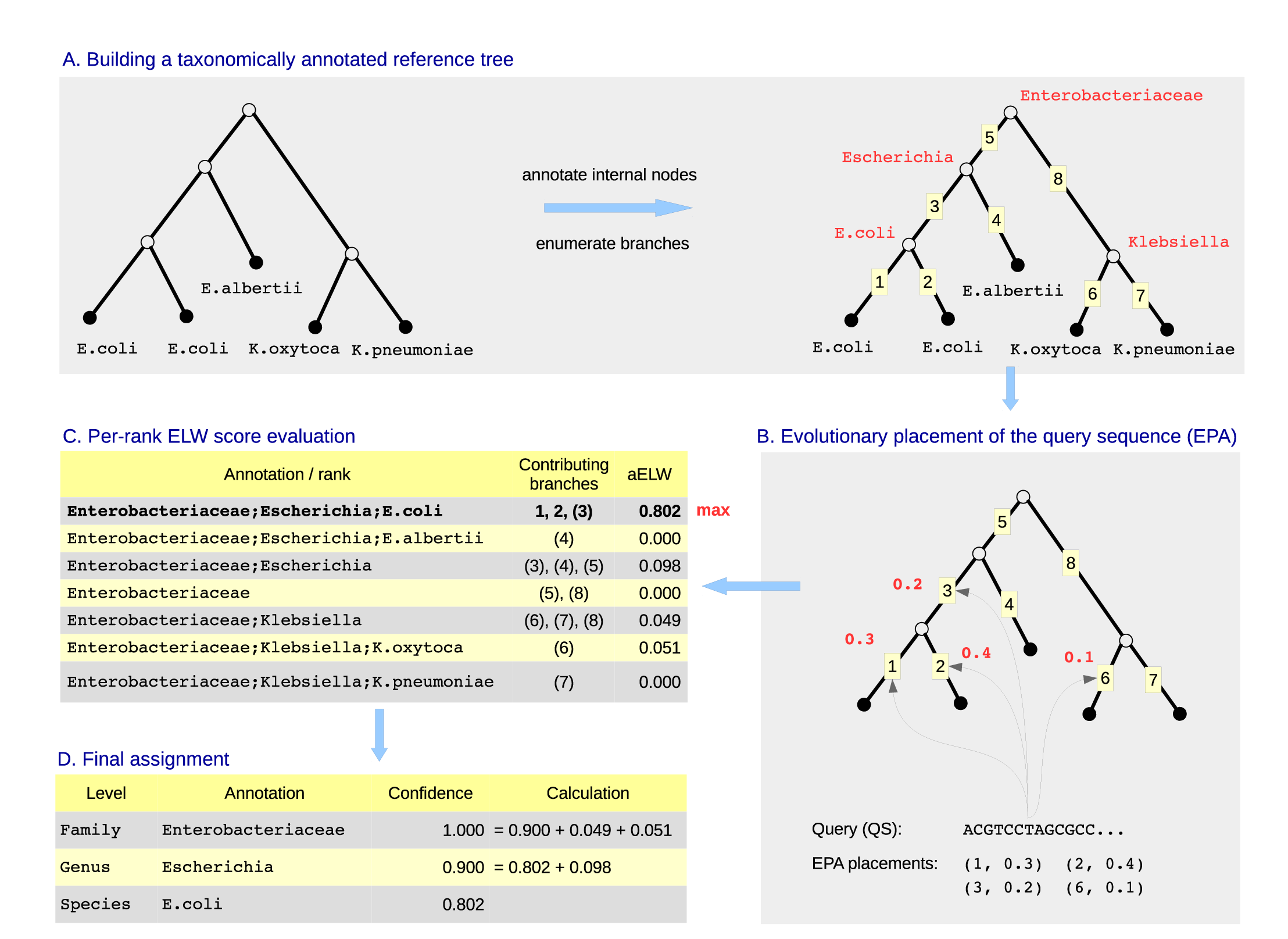
Example illustrating the taxonomic assignment method implemented in SATIVA. **A.** We start with a resolved, bifurcating reference tree which has taxonomic annotations at the tips (i.e., sequence annotations). First, we perform a post-order tree traversal and assign taxonomic labels to internal nodes (shown in red) by taking the longest common annotation of the respective child nodes. Then, we enumerate the branches and store the mapping between the branch numbers and the taxonomic annotations of their adjacent nodes. **B.** We use the RAxML-EPA algorithm to obtain the most likely placements of the QS on the reference tree. For our purposes, each placement is represented by a pair (branch number, likelihood weight). **C.** We compute the accumulated likelihood weight (aELW) for each taxonomic rank by summing over the weights of the corresponding branches. The branches in parentheses have two competing annotations, and their weights contribute partially to both respective aELWs (see main text for details). **D.** We assign the QS to the taxonomic rank with the highest aELW. At each taxonomic level, we compute a confidence score by summarizing aELWs of all annotations which do not contradict the assigned one at this level.

#### 2) *Taxonomic assignment.:*

We use the following approach to assign a taxonomic annotation to a so-called *query* sequence (QS), that is, a sequence which is not present in the reference tree. First, we use the Evolutionary Placement Algorithm (EPA, as implemented in RAxML) to calculate the most likely placement(s) of the query in the reference tree (see Fig. 2, B). In particular, for each branch of the reference tree we obtain a *expected likelihood weight* (*ELW*) value (24; 25). The ELW is calculated as the ratio of the likelihood of the tree including the QS placed into the current branch *b*_*k*_ to the sum of likelihoods for all possible QS placements:

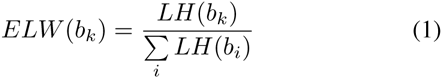

Once all likelihood weights for a QS have been computed, we use them to calculate the accumulated ELW (aELW) for each taxonomic rank (see Fig. 2, C). In order to map branches to taxonomic ranks, we make use of the taxonomically-labeled reference tree constructed in the previous step. In particular, for each branch *b*_*k*_, we analyze the taxonomic annotations *r*_*u*_ and *r*_*l*_ of the nodes *adjacent* to this branch. We distinguish two cases:

(a) If *both* taxonomic annotations are identical (*r*_*u*_ = *r*_*l*_ = *r*_*i*_), then the entire likelihood weight of the branch will contribute to the aELW of this annotation:

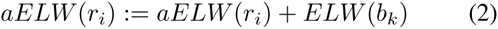
(b) If the taxonomic annotations differ, then the likelihood weight is distributed between both annotations:

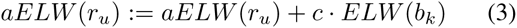

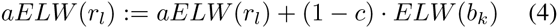

where parameter *c* defines the distribution ratio. In the current implementation, we distribute weights (almost) equally by setting *c* := 0.49 (a small imbalance is needed to avoid ties).

By applying this procedure to all branches with the placements, we obtain a single aELW score for each taxonomic annotation. Thereafter, we select the taxonomic annotation with the highest aELW as new, phylogeny-aware annotation for the QS.

Note that ‘nested’ taxonomic annotations such as (*’Escherichia’, ‘E.coli’*) and (*’Escherichia’*) are considered as distinct at this step of our algorithm. This allows to directly compare competing annotations at *different* taxonomic levels to each other and select the most likely one according to the aELW.

Finally, we calculate an overall *assignment confidence score* for the QS by summing over the aELWs for *all* annotations that are in concordance with the *proposed* taxonomic annotation of the QS. So, if (*’Enterobacteriaceae’, ‘Escherichia’*) is selected as the most likely QS assignment, the aELW for the annotation (*’Enterobacteriaceae’, ‘Escherichia’, ‘E.coli’*) will also contribute to the assignment confidence score at both, the family and genus level. At the same time, aELWs for (*’Enterobacteriaceae’, ‘Klebsiella’*) or (*’Enterobacteriaceae’*) annotations will contribute to the family assignment confidence score, but not to the genus level assignment confidence score (see Fig. 2, D).

#### 3) Identification of mislabels.

The mislabel identification process consists of two steps:

1) **Leave-one-out test.** We prune *one sequence at a time* from the reference tree, and use EPA to place it back into all branches of the remaining reference tree. Then, we use the taxonomy assignment approach described above to calculate a new taxonomic label for the pruned sequence. If there is a disagreement between the new and the original taxonomic label, we put the sequence into the *preliminary* mislabels list. Note that, when comparing taxonomic labels, we do not consider missing rank annotations as disagreements. For instance, given that the original annotation is (*’Enterobacteriaceae’, ‘Escherichia’),* new annotations (*'Enterobacteriaceae'*, *'Klebsiella'*) or (*’Enterobacteriaceae’, ‘Klebsiella’, ‘K.pneumoniae’*) will be reported as a mislabel, whereas (*'Enterobacteriaceae'*, *’Escherichia’, ‘E.coli’*) and (*’Enterobacteriaceae’*) will not.
2) **EPA test.** Once all sequences have been subjected to the leave-one-out test, we now prune all sequences in the preliminary mislabels list from the reference tree *at once*. Then, we use EPA to independently place each of them back into the remaining tree and re-calculate the annotations. Once again, we compare the new annotations with the original ones, and put sequences in the *final* mislabels list if they differ. In addition, the calculated taxonomic annotation for each final mislabel sequence will be reported as a suggested correction. Further, we report the *mislabel confidence score*, which is equal to the assignment confidence score (see above) at the *highest* taxonomic rank level for which the original and the new taxonomic labels differ.

Although we could have used a single leave-one-out test to identify mislabels, our initial experiments showed that this yielded a higher false positive rate. We assume that this is due to the noise introduced by the mislabels in the reference phylogeny. Note that, in this first step, all but one mislabel are still contained in the reference phylogeny. In particular, we observed a phenomenon which we call ‘reciprocal’ mislabels: if one out of two highly similar (’buddy’) sequences was taxonomically mislabeled (and thus placed in a remote clade of the reference tree), both of them were incorrectly reported as mislabels. For each sequence, a re-classification into the rank of its buddy was proposed in the leave-one-out test. In other words, suggested corrections were reciprocal. This situation can be resolved by simultaneously pruning both sequences from the reference tree and placing them back independently. Therefore, we use the two-step approach presented above.

#### 4) Implementation details and ARB integration.

The tool is implemented in a Python pipeline, which calls the RAxML executable to calculate the taxonomically constrained reference phylogeny and to perform evolutionary placements (see Fig. 1). The remaining workflow steps (e.g., branch labeling and taxonomic assignment) are implemented in Python using the ETE library (26). The framework requires two input files: a multiple sequence alignment (FASTA or PHYLIP) and a list of taxonomic annotations with matching sequence identifiers. The output is a tab-delimited text file which contains the putative mislabel identifiers as well as the original and suggested taxonomic annotations including confidence values.

To make the tool easy-to-use for taxonomists, we integrated SATIVA with the ARB software (27), which is a widely used tool for maintaining and curating large rRNA databases. ARB is built around an efficient sequence storage engine, it provides a GUI for editing sequences and associated metadata as well as import/export tools, and offers advanced tree visualization capabilities. SATIVA is currently available in the development version of ARB, which can be downloaded at http://www.arb-home.de/downloads.html. Within ARB, the user can invoke SATIVA by simply marking a subset of sequences in the GUI and selecting “Validate taxonomy” from the menu. The results of the analysis are written back to the ARB database fields, and putatively mislabeled sequences will be highlighted on the tree (see Fig. 3). ARB can be used to easily visualize results and customize the visualization using ARB features such as “search by field value” and “set field value”. For instance, mislabeled sequences can be highlighted with different colors based on their rank level incongruence (e.g., phylum, class etc.) or mislabel confidence value. Finally, ARB can also write SATIVA results to an external file.

**Fig. 3:**
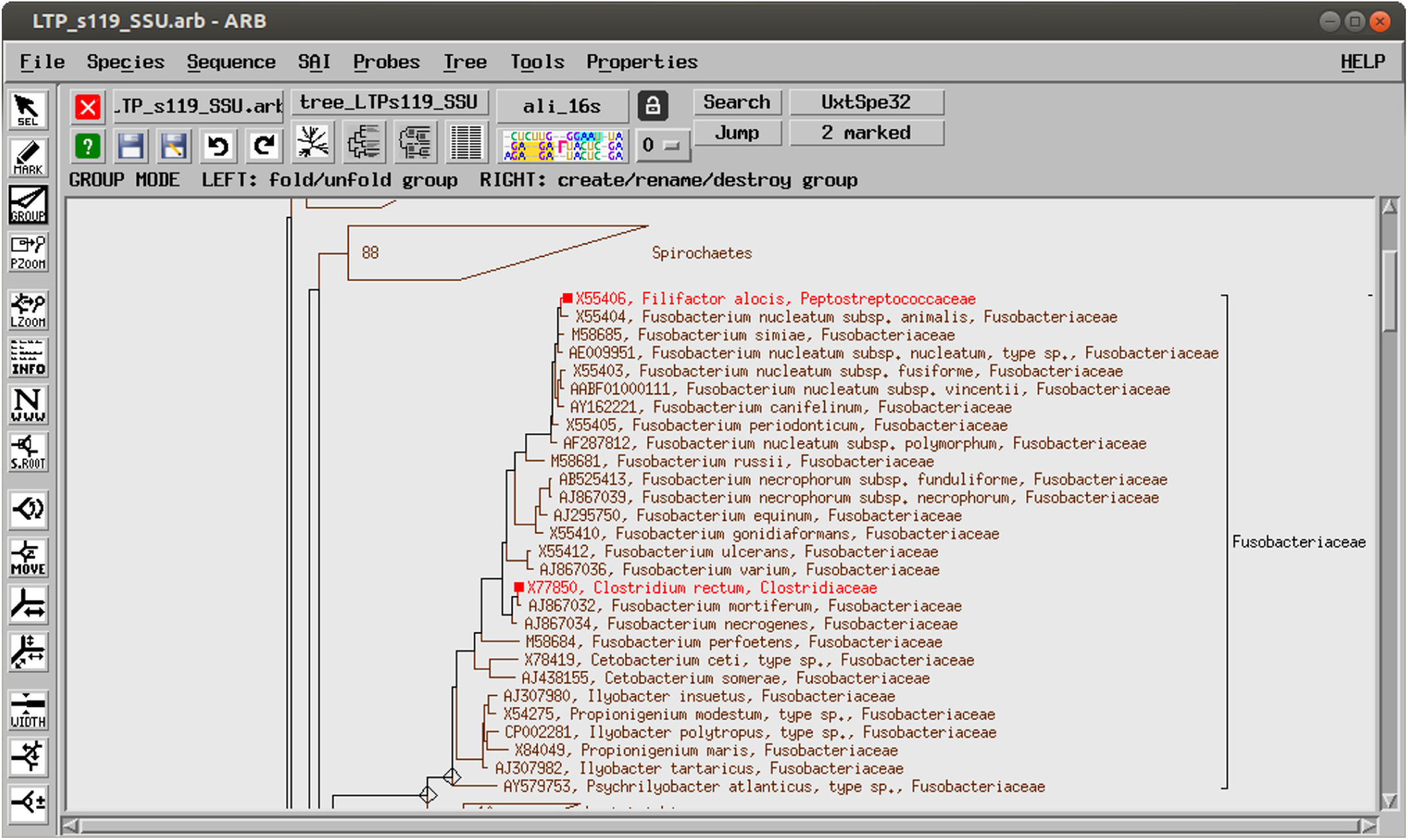
SATIVA results displayed in ARB. Two sequences annotated as *Clostridiales* species in LTP123 have been identified as mislabels (marked in red). The suggested re-classification as *Fusobacterium* is consistent with the SILVA123 annotation (placement in the tree).

The standalone as well as ARB-integrated versions of SATIVA provide parameters which allow to set the trade-off between runtime and the thoroughness of mislabel detection. SATIVA can perform multiple RAxML tree searches (with different starting trees) to find the best-scoring reference tree. The number of tree searches can be set by the user and affects the runtime linearly. SATIVA can also be run in the ‘fast’ mode, in which a topological convergence criterion is used to stop the tree search earlier (corresponds to ‐D option of RAxML (28)). Conversely, in the ‘thorough’ mode (default) SATIVA uses a likelihood-based stopping criterion, which is typically 1.5 to 2.1 times slower (28).

### B Alternative methods

To the best of our knowledge, there are no established tools for automatic mislabel identification. Thus, a direct accuracy and performance comparison is not feasible at present. However, since with SATIVA we introduce a novel method for taxonomic assignment of new sequences, we included two widely used taxonomic classification methods (UCLUST (29) and RDP (30)) into our performance evaluation. To perform an as fair as possible comparison, we emulated a leave-one-out approach for assessing both tools, which is analogous to the one implemented in SATIVA. For this, we remove one sequence at a time from a database with *n* sequences and use the remaining sequences and taxonomic labels as new reference. Then, we use RDP and UCLUST to assign a new taxonomic label for the removed sequence using the new reference with *n* − 1 sequences. If the new taxonomic label is different from the existing one, we consider it as mislabel and the inferred taxonomic label as the proposed new classification. For both RDP and UCLUST, we used the implementations that are available in the QIIME v1.8.0 (31)pipeline with default parameters.

### C. Simulated data

In order to test the performance of different mislabel identification approaches, we generated simulated trees and datasets. We superimposed a 'true' taxonomic classification that is fully congruent with the phylogeny used to simulate the data (for further details, see below). We then randomly mislabeled a small fraction of the sequences at each taxonomic rank (for details, see below), and subsequently tested if the algorithms can identify these mislabels and suggest the correct rank.

We used the LTP (version 123) dataset as the basis for our simulations. First, we generated a ML-based fully bifurcating tree with RAxML that uses the LTP taxonomy as constraint tree and the LTP reference alignment as input. Next, we partitioned the alignment in order to reflect the known secondary structure of the 16S rRNA gene: we defined nine partitions for variable regions V1-V9, one for all conserved regions, and one for the ‘flanking’ regions not found in the reference *E.coli* sequence. We estimated the GTR (General Time Reversible substitution matrix) parameters and branch lengths for each partition individually. These parameters as well as the ML tree topology were subsequently used to simulate sequence alignments with INDELible (32). We tuned the INDELible simulation parameters (insertion/deletion rate and sequence length at the root) such that for each partition, the length and proportion of gaps approximately match the corresponding region of the empirical alignment.

Finally, we set the target overall percentage of mislabels as well as an error probability at each taxonomic rank (e.g., 1% of mislabels in total, of which 5% have an incorrect phylum, 10% an incorrect class etc.). Then, we randomly selected a subset of sequences and substituted their original labels with different (“incorrect”) ones according to the percentages defined above. We generated two simulated datasets, which have 1% and 5% mislabels, respectively. For each mislabel rate, we generated 3 replicates with different random number seeds.

Further details on simulation are available in the Supplementary Material. Scripts and data files can be downloaded from https://github.com/amkozlov/mislabels16-data.

### D. Real-world datasets

We analyzed four established databases of taxonomically annotated 16S rRNA sequences of Bacteria and Archaea: RDP-II (15), Greengenes (16), the ‘All-Species’ Living Tree Project (LTP) (33), and SILVA (14). These databases have different underlying taxonomies, and vary in their size and taxonomic composition. In particular, LTP is a highly curated 16S and 23S rRNA gene sequence database, which includes bacterial and archaeal type strains only, and thus has a moderate number of sequences (11,939 as of release 123, September 2015). RDP-II, Greengenes, and SILVA, however, contain all rRNA sequences available in public databases that passed a quality check. Hence, they contain orders of magnitude more entries (1.2–3 million sequences). Furthermore, SILVA and Greengenes provide non-repetitive (NR) subsets of sequences clustered with a 99% identity threshold, referred to as ‘NR99’ henceforth.

In order to make results for different taxonomies comparable and to maximize the coverage of each individual database, we divided our analysis into two parts. First, we evaluate taxonomic annotations of type strains only, using the same sequence set and alignment (LTP v123) for all four databases (datatsets `GG13_T`, `LTP123_T`, `RDP11_T` and `SLV123_T` in Table 1). Second, we evaluate the NR99 subsets (that is, representatives of the 99% identity clusters) for Greengenes and SILVA (datasets `GG13_NR99` and `SLV123_NR99`), thereby also including environmental sequences into our analysis.

**Table 1.**
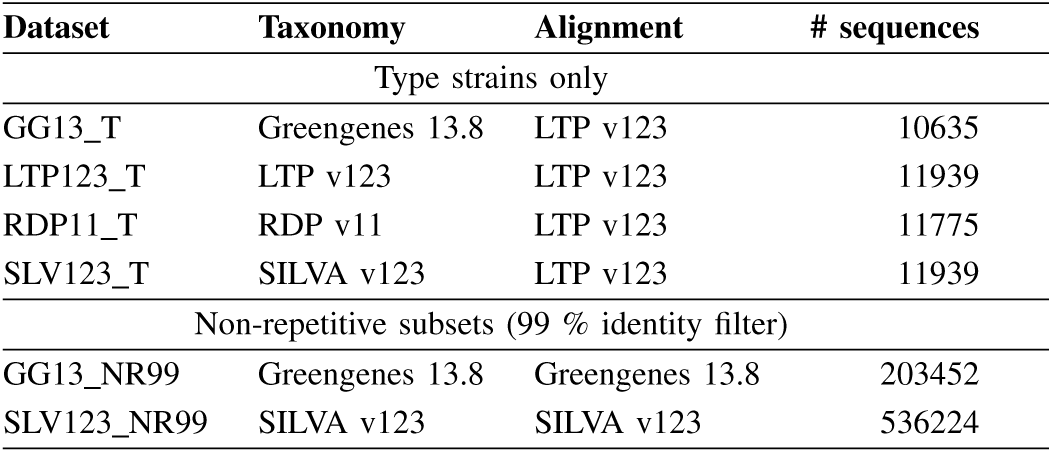
Real microbial 16S datasets

| Dataset | Taxonomy | Alignment | # sequences |
| --- | --- | --- | --- |
| Type strains only |  |  |  |
| GG13_T | Greengenes 13.8 | LTP v123 | 10635 |
| LTP123_T | LTP v123 | LTP v123 | 11939 |
| RDP11_T | RDP v11 | LTP v123 | 11775 |
| SLV123_T | SILVA v123 | LTP v123 | 11939 |
| Non-repetitive subsets (99 % identity filter) |  |  |  |
| GG13_NR99 | Greengenes 13.8 | Greengenes 13.8 | 203452 |
| SLV123_NR99 | SILVA v123 | SILVA v123 | 536224 |

For the type strain datasets, we executed SATIVA in ‘thorough’ mode, using 10 RAxML runs to infer the reference tree. For the NR99 datasets, we used ‘fast’ mode and 1 RAxML run, for computational reasons. The confidence cut-off was set 0.51 for all datasets.

Further details on empirical dataset assembly are available in the Supplementary Material.

### E. Case study: Cyanobacteria

In addition to the simulated and full Bacteria/Archaea datasets, we also examined a single bacterial phylum, the *Cyanobacteria*, more extensively. This phylum is known to be taxonomically challenging, and several alternative taxonomic frameworks have been proposed to classify *Cyanobacteria* (34).

With this case study we intend to assess to which extent SATIVA is useful in practice for evaluating and improving ‘difficult’ taxonomies.

First, we analyzed four existing taxonomies: European Nucleotide Archive/GenBank (EMBL), SILVA, Greengenes, and RDP-II. Then, we used the “Candidate Taxonomic Unit” (CTU) recognition process as proposed by Yarza *et al*. (35) to construct a novel, unified taxonomic framework for the cyanobacterial sequences (called CyanoCTU henceforth). The CTU method offers a simple procedure to devise phylogeny-aware taxonomies by overlaying a phylogenetic tree with OTU (Operational Taxonomic Unit) information.

To infer a phylogenetic tree, we used an alignment of 1050 quality-checked, full-length (>1400 bases) 16S rRNA sequences from the SILVA SSU Ref database (selected based on the organism name and strain metadata field). This alignment was produced by SINA (36) and is available from SILVA. We filtered this alignment with a 10% cyanobacterial base conservation threshold. Then, we ran RAxML (v. 7.7.2 (37)) to generate 1000 rapid bootstrap replicates and performed a subsequent search for the best-scoring ML tree under the GTR+Γ model.

To generate OTUs, we performed a hierarchical furthest neighbor clustering using MOTHUR v1.20.3 (38) with specific sequence identity thresholds for each taxonomic rank level (75% phylum, 78.5% class, 82% order, 86.5% family, and 94.5% genus (35)). Then, we applied a custom `Perl` script (Supplementary file 3) to assign each sequence to five distinct OTUs, one per taxonomic rank level. This information was imported into ARB (27), such that each sequence in the phylogeny is annotated with the assigned OTU labels.

In the last step, we used the ARB tree editing functions to manually assign taxonomic rank names to monophyletic clades on the phylogenetic tree, taking into account both, the tree topology and the OTU labels of individual sequences. For naming taxonomic ranks, we used the nomenclature that follows the format `Rank_NameN-M`. Here, `Rank_name` is simply the name of the taxonomic rank level, that is, ‘Genus’, ‘Family’, ‘Order’, ‘Class’, or ‘Phylum’. The number N is used as an identifier for an OTU at a particular taxonomic rank level. For instance, *Class10* denotes the 10th OTU found at the class level or *Genus126* the 126th OTU found at the genus level. M is an optional number used to discriminate between multiple taxonomic ranks derived from one and the same OTU if this OTU is polyphyletic (e.g., *Family5-1* or *Family5-2*).

Finally, we applied SATIVA to evaluate the phylogenetic consistency of the novel CyanoCTU taxonomy and compare it to the existing taxonomies mentioned above. SATIVA was run with the following parameters: thorough inference, 10 RAxML searches, and a confidence cut-off of 0.51. Out of the 1050 cyanobacterial sequences used in phylogenetic inference, only 900 were used, as 150 sequences were listed as ‘Unclassified’ by either RDP-II or Greengenes. Species level mislabels were only considered for EMBL taxonomy, as this is the only taxonomy where species names are consistently always present.

## III. RESULTS

### A. Performance on simulated data

We deployed two metrics to quantify the ability of competing tools to identify mislabels on simulated data. Firstly, we used the accuracy of mislabel *identification*. To this end, we compared the output of each program to the true list of mislabels: each sequence was counted as *true positive* (*TP*) if it was present in both lists, and as *false negative (FN)* or *false positive (FP)* if it was missing from the inferred or ground truth list, respectively. Then, we used the standard formulas to calculate *precision* and *recall* values at each taxonomic level (s. Table 2). Secondly, we evaluated the *correction accuracy* by comparing the suggested taxonomic annotation for mislabels with the true one (s. Table 3). If a mislabel was not identified as such, we assumed its inferred annotation to be equal to the original, uncorrected one. In other words, such sequences were counted as *false positives* at taxonomic levels that were (deliberately) mislabeled in the respective simulations.

**Table 2.** Accuracy of mislabel identification. Taxonomic levels are the levels where sequences were deliberatly misclassified in the ground truth. That is, a recall value of 0.974 at the family level means that 97.4 % of sequences with an incorrect family label were successfully identified.

| Level | Percentage of mislabeled sequences |  |  |  |  |  |  |  |  |  |  |  |
| --- | --- | --- | --- | --- | --- | --- | --- | --- | --- | --- | --- | --- |
|  | 1 % |  |  |  |  |  | 5 % |  |  |  |  |  |
|  | Precision |  |  | Recall |  |  | Precision |  |  | Recall |  |  |
|  | RDP | UCLUST | SATIVA | RDP | UCLUST | SATIVA | RDP | UCLUST | SATIVA | RDP | UCLUST | SATIVA |
| Phylum | 0.625 | 1.000 | 1.000 | 1.000 | 0.933 | 1.000 | 0.813 | 0.947 | 1.000 | 1.000 | 0.828 | 1.000 |
| Class | 0.675 | 0.958 | 0.900 | 1.000 | 0.852 | 1.000 | 0.785 | 0.896 | 0.983 | 0.989 | 0.793 | 1.000 |
| Order | 0.409 | 0.796 | 1.000 | 1.000 | 0.867 | 1.000 | 0.661 | 0.860 | 0.977 | 0.995 | 0.726 | 0.995 |
| Family | 0.311 | 0.657 | 0.965 | 0.973 | 0.811 | 0.991 | 0.605 | 0.833 | 0.971 | 0.976 | 0.745 | 0.987 |
| Genus | 0.054 | 0.130 | 0.893 | 0.965 | 0.832 | 0.965 | 0.217 | 0.416 | 0.822 | 0.949 | 0.757 | 0.928 |
| Total | 0.120 | 0.274 | 0.939 | 0.977 | 0.836 | 0.984 | 0.382 | 0.619 | 0.917 | 0.971 | 0.757 | 0.969 |

**Table 3.**
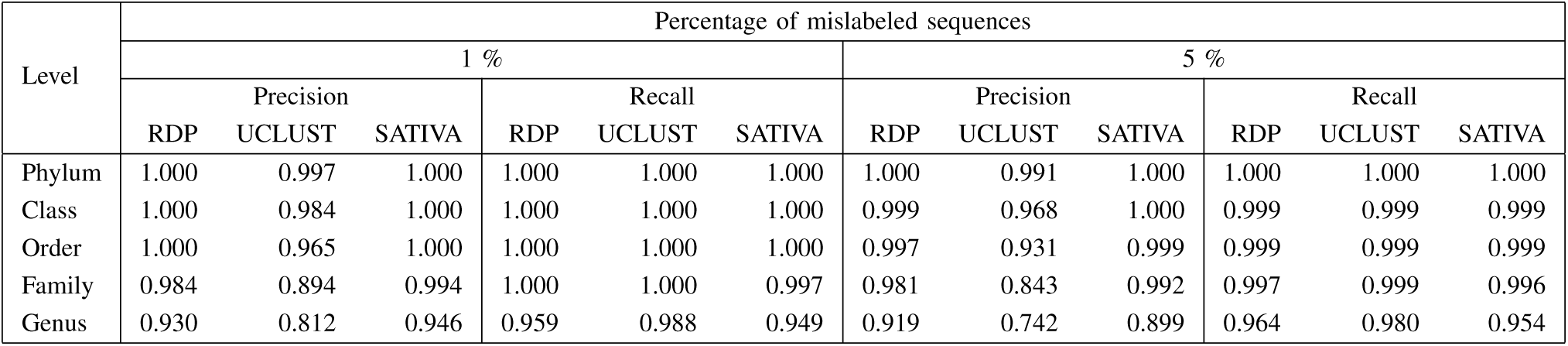
Accuracy of the suggested taxonomic annotation for mislabels. Note that, errors are propagated down the taxonomy,that is, an incorrect family label also implies an incorrect genus label, etc.

Since all three methods provide confidence values for taxonomic placements (RDP, UCLUST) or identified mislabels (SATIVA), it is possible to use a threshold to exclude results with low confidence. For each method, we empirically evaluated several confidence thresholds and chose the value which yielded the highest F-measure value (that is, the best precision/recall trade-off). Specifically, we set the confidence threshold to 0.7 for UCLUST, 0.8 for RDP and 0.51 for SATIVA.

Among the three algorithms tested, SATIVA shows the best mislabel identification accuracy: at least 96.9% of all mislabels with wrong annotations are recognized, while the false positive rate is less than 9%. The RDP classifier achieved similar recall values to SATIVA (e.g., 97.7% vs. 98.4% on the dataset with 1% mislabels). However, its precision was unacceptably low (12.0% / 38.2%). Finally, the UCLUST algorithm shows higher precision, but lower recall than RDP, and is clearly inferior to SATIVA in terms of both precision and recall.

Our measurements of *precision* for UCLUST and RDP might appear contradictory to earlier studies (e.g., (30)), where much higher values have been reported. Note that, here we measure the precision of *mislabel identification*, which is different from the precision of *taxonomic classification*. Specifically, in the latter case *all* sequences with correctly inferred taxonomic annotation are considered *true positives*. In our test, however, only those sequences that were *deliberately mislabeled* and correctly identified by the method are counted as *true positives*. All other, non-mislabeled sequences, which were recognized as such, represent *true negatives*. And since in our test datasets mislabeled sequences constitute only a small fraction of the data (1% or 5%), the impact of *false positives* on precision is much more pronounced. This also explains the significantly higher precision values for the 5% dataset as compared to the 1% dataset.

In the correction accuracy test, SATIVA and RDP performed almost equally well, achieving precision and recall of around 95% (although precision drops to ≈90% on the dataset with 5% mislabels for both methods). UCLUST showed higher recall (98.8% / 98.0%), but this comes at the expense of a substantially lower precision (81.2% / 74.2%).

### B. Retrospective assessment of established 16S taxonomies

We used our approach to assess the phylogenetic consistency and identify mislabels in four widely-used 16S databases. We ran SATIVA on the representative datasets (see Section Real-world datasets for details) and evaluated the percentage of mislabels reported for each taxonomic rank (see Fig. 4) as well as for several major bacterial phyla (s. Fig. 5).

**Fig. 4:**
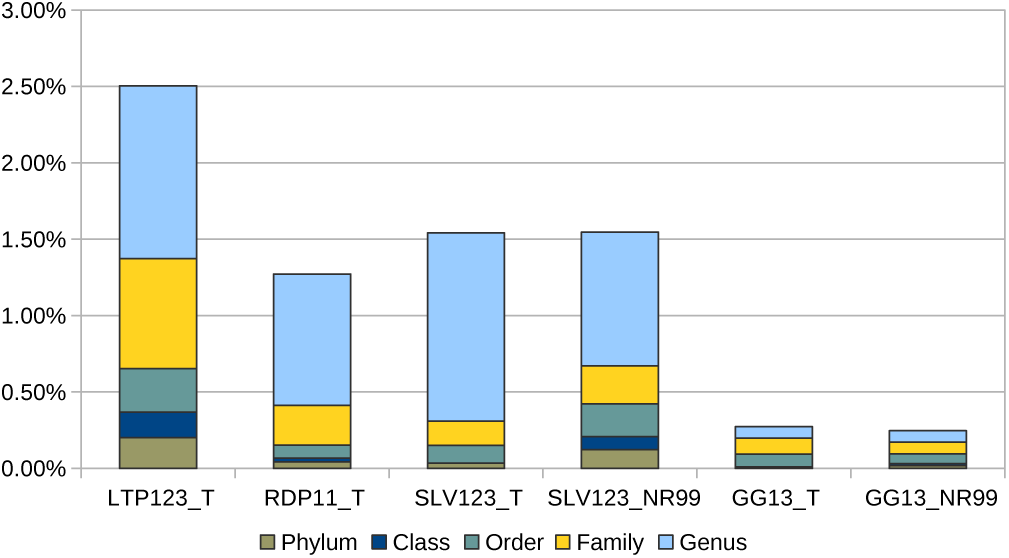
Percentage of mislabels by rank (cumulative) for widely-used 16S taxonomies.

**Fig. 5:**
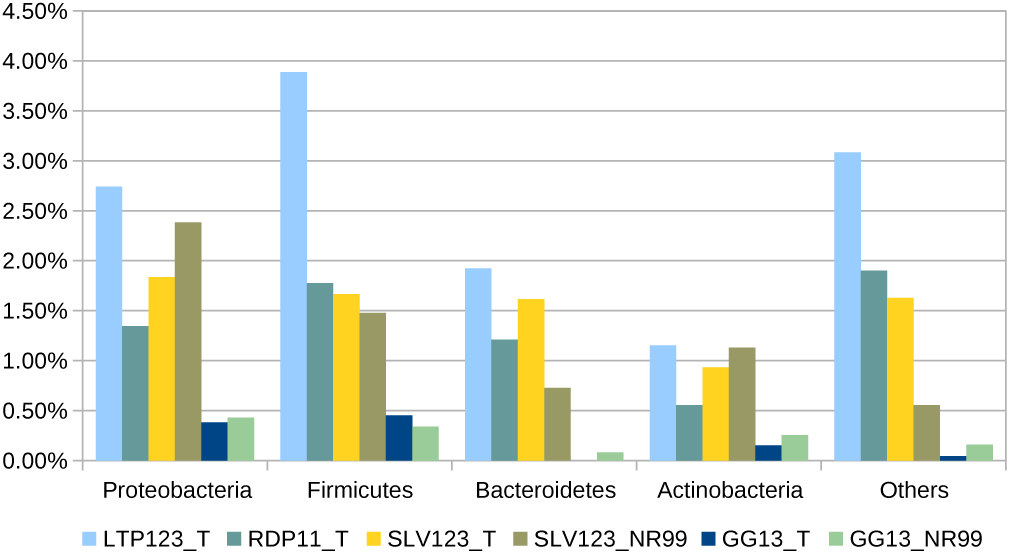
Percentage of per-phylum mislabels for widely-used 16S taxonomies.

Among type strain datasets, `GG13_T` exhibits by far lowest percentage of identified mislabels (0.27%), followed by `RDP11_T` (1.27%), `SLV123_T` (1.54%) and `LTP123_T` (2.52%). In all taxonomies but Greengenes, the vast majority of mislabels was detected at the genus level. Therefore, the estimated percentage of mislabels at higher taxomomic levels (family and above) is more similar across datasets: `GG13_T` 0.20%, `SLV123_T` 0.31%, `RDP11_T` 0.41% and `LTP23_T` 1.37%.

On the datasets which also include environmental sequences (`GG13_NR99` and `SLV123_NR99`), the Greengenes taxonomy shows less inconsistency (0.27% mislabels) compared to SILVA (1.55%). Again, the difference becomes less pronounced if genus-level mislabels are excluded (0.17% vs. 0.67%).

As Fig. 5 shows, the distribution of mislabels among individual phyla is non-uniform. In all taxonomies, *Actinobacteria* and *Bacteroidetes* appears to contain less mislabels (0.15%–1.15% and 0%–1.92%, respectively) than *Proteobacteria* (0.38%-2.74%) and *Firmicutes* (0.34%-3.89%).

The complete lists of identified mislabels for each 16S taxonomy are provided in the Supplementary File 1.

### C. Cyanobacteria taxonomy

The EMBL taxonomy for *Cyanobacteria* recognizes six orders as *Chroococcales*, *Nostocales*, *Oscillatoriales*, *Pleurocapsales, Prochlorales*, and *Stigonematales*. SILVA recognizes five subsections, which could be considered as orders, while RDP-II has thirteen families. The Greengenes taxonomy is more similar to the EMBL taxonomy, in the sense that, it also recognizes six orders as *Chroococcales*, *Nostocales*, *Oscillatoriales*, *Pseudanabaenales*, *Stigonematales*, and *Synechococcales*. These different taxonomic groups are not necessarily congruent among the different taxonomies, as shown by the superimposition with our reference phylogeny (see Section Case study: Cyanobacteria and Supplementary Figure S1). Most groups, except for some smaller groups in the RDP-II taxonomy, are polyphyletic. For the EMBL and SILVA taxonomies, certain groups do overlap.

SATIVA runs on these reference taxonomies for Cyanobacteria returned between 0% and 11.5% mislabels. Specifically, 104 mislabeled sequences (11.5%) were detected for the EMBL taxonomy, 9 (1%) for SILVA, 0 for RDP-II, and 1 (0.1%) for Greengenes. In all taxonomies, most mislabels were identified at the genus level. The mislabels were distributed across all orders in the EMBL taxonomy, but were restricted to *Nostocales* in Greengenes taxonomy, and *FamilyI* in SILVA taxonomy.

As explained above (see Case study: Cyanobacteria), the CyanoCTU taxonomy was built using hierarchical clustering. In particular, we obtained one class-level OTU, two order-level OTUs, fourteen family-level OTUs, and 171 genus-level OTUs after clustering. These OTUs were further refined manually to form taxonomic ranks. Specifically, we decided to merge some OTUs together based on additional evidence such as the phylogenetic tree topology, distances between the sequences (calculated with the `arbdist` tool), or expert judgement. Finally, we included one class, one order, five families, and 126 genera in our novel CyanoCTU taxonomy. SATIVA reported only 2 mislabels for CyanoCTU, both at genus level.

Full lists of identified mislabels for all evaluated *Cyanobacteria* taxonomies are provided in the Supplementary File 2.

### D. *Memory consumption and running time*

Computational complexity and memory requirements of SATIVA are dominated by the phylogenetic methods it employs, and their specific implementation in RAxML. The additional overhead induced by the remaining steps of the pipeline is negligible.

Memory requirements of SATIVA are determined by the EPA: due to its parallelization scheme (39), EPA memory footprint is approximately three times higher compared to standard tree inference. SATIVA memory consumption for a specific dataset can be easily estimated using the RAxML online calculator available under http://sco.h-its.org/exelixis/web/software/raxml/index.html#memcalc.

The two largest contributions to run time are: the topologically constrained ML tree search (to resolve multifurcations in the taxonomic constraint tree) and the leave-one-out test. Apart from the obvious dependence on alignment dimensions, tree search run times are influenced by the degree of resolution of the taxonomic constraint and also by the phylogenetic signal in the alignment. As for ML tree inference in general, run time prediction for this step is difficult. In contrast to this, the computational complexity of the leave-one-out test depends solely on the dataset dimensions: namely, alignment width (linear) and number of sequences (quadratic, but can be reduced to quasi-linear using the EPA heuristics (22), which is the default algorithm used by SATIVA for datasets with more than 1,000 sequences).

The real-world datasets analyzed in this study differ greatly in their dimensions, in terms of both number of taxa and alignment width. Hence, we chose experimental platform configuration according to computational requirements and parallelization potential of each particular dataset.

We conducted the analysis of the small *Cyanobacteria* datasets (≈ 1000 taxa) on an 8-core server, where it took about 1 hour to complete. For the type strain datasets (≈ 10,000 taxa), we used a single 28-core node of the SuperMUC super-computer. The running time was between 34 hours (`GG13_T`) and 47 hours (`LTP123_T`). For the largest NR99 datasets, we used a HITS cluster node equipped with 32 CPU cores and 512GB of RAM (reference tree inference) as well as multiple nodes of the SuperMUC (leave-one-out test). In this configuration, the overall runtime was ≈ 100 hours for the `GG13_NR99` dataset and ≈ 480 hours for the SLV123_NR99 dataset.

In all experiments, SATIVA was executed with 1 thread per physical CPU core (default setting). Additional details on individual hardware configurations and running times are provided in the Supplementary methods.

## IV. DISCUSSION

### A. Algorithm rationale and comparison to other approaches

As already mentioned, SATIVA performs an EPA-based leave-one-out test based to detect inconsistencies between taxonomic annotations and phylogenetic signal in the alignment. It was previously shown, that EPA can reliably re-insert a pruned sequence into its original position on a best-known maximum likelihood (ML) tree (22). In our case, however, the reference phylogeny is not the best-known ML tree, since a potentially erroneous (inconsistent with the phylogenetic signal in the data) taxonomy was used to constrain the tree inference process. Intuitively, if the proportion of mislabels is low (i.e., there is little conflict between taxonomy and phylogeny), then, phylogenies obtained via a constrained and unconstrained search will be mostly congruent. Consequently, EPA placement accuracy on the taxonomically constrained reference tree will be almost as high as on the best-known ML tree. This hypothesis is confirmed by our simulation results: SATIVA shows high accuracy on simulated datasets with 1% and 5% mislabels.

As we show in simulation, SATIVA is superior to leave-one-out tests relying on the RDP Classifier or UCLUST with respect to mislabels identification. Additionally, the RDP Classifier needs to be re-trained from scratch after pruning each sequence. This makes it computationally inefficient even on medium-size datasets. In SATIVA, we circumvent this retraining overhead by assuming, that pruning a single taxon will not induce substantial changes into the reference phylogeny. So, instead of conducting a *de novo* tree inference, we simply re-calculate the likelihood of the remaining reference tree with one sequence less. Finally, the UCLUST method, which is based on fast alignment heuristic implemented in USEARCH (29), does not require an explicit training step and was approximately as fast as SATIVA in our experiments.

### B. Mislabels in established 16S taxonomies

According to our analysis, Greengenes, RDP, SILVA, and, to a lesser extent, LTP, are consistent at higher taxonomic levels. The few sequences proposed for re-classification into a different phylum or class are most probably due to an incorrect culture in the collection. Although putatively mislabeled sequences are more common at lower ranks (e.g., family or genus), their overall percentage is below 3% for all taxonomies. This implies that current taxonomic frameworks represent the phylogenetic signal of 16S rRNA well, but that there is nevertheless room for improvement.

We identified the highest amount of putative mislabels in the LTP taxonomy, especially at higher taxonomic ranks. This can be explained by the fact, that the LTP classification strictly follows Bergey’s taxonomic outlines and LPSN. Conversely, the other three taxonomies adapt their classifications in order to better reflect the 16S tree topology, even if it involves changes that violate the formal rules of taxonomic and nomenclature code. For instance, non validly published names are widely used to split non-monophyletic taxa (e.g., *’Clostridium III’* or *’Bacillaceae 1’)* or to represent uncertainty in classification (e.g., *’Clostridiales Incertae Sedis’*). Although such changes might be justified from the practical standpoint, they make direct comparison between LTP and other taxonomies impossible. Therefore, we suggest that LTP should be viewed as a ‘baseline’ in our comparison.

At the other end of the scale, the Greengenes taxonomy shows an extremely low percentage of mislabeled sequences (<0.3%). This suggests that this taxonomy is phylogenetically very consistent, most likely owing to the fact that it is based on a de novo phylogeny. On the other hand, the anomalously few genus-level mislabels (0.08%) could be partially explained by the lack of annotation at this level. More specifically, as much as 29% of the sequences in `GG13_T` and 54% in `GG13_NR99` are not assigned to a genus. For comparison: just 0.04% of the sequences in `SLV123_T` and 16% in `SLV123_NR99` do not have a genus-level annotation.

Interestingly, the overall percentages of identified mislabels in full non-repetitive datasets (GG13_NR99 and SLV123_NR99) and in the corresponding type strain datasets (GG13_T and SLV123_T) are highly similar. However, SILVA_NR99 shows substantially more mislabels at higher taxonomic levels compared to SLV123_T (0.12% vs. 0.03% at the phylum level and 0.08% vs. 0% at the class level). This finding suggests that in the SILVA database, at least the most obvious misannotations were fixed for type strain sequences.

### C. Cyanobacteria taxonomy

Cyanobacteria were chosen specifically as this phylum represents a classification challenge, due to the dual nomenclature code employed – the botanical and bacteriological code (40), and due to numerous classification schemes being in place. Komárek *et al*. have recently provided a comprehensive review of the current cyanobacterial classification systems (34). Additionally, the authors have also proposed a new classification based on a molecular phylogeny of 31 conserved proteins.

Given that the different groups each taxonomic hierarchy recognizes are by no means monophyletic, the number of mislabels found is surprisingly low, indicating that at least each group is consistently defined. In case of EMBL, the identified mislabels were most likely a result of incorrect organism names in the original sequence records. Additionally, in case of the SILVA taxonomy, some sequences obviously had a suboptimal placement in the original guide tree, thus leading to misclassifications.

As we identified few (or no) mislabels in SILVA, RDP-II, and Greengenes taxonomies, all of them can be considered usable for Cyanobacteria. However, a taxonomic framework ideally should aspire to have monophyletic taxa, and none of the frameworks considered here satisfy this condition. Of the different taxonomic hierarchies considered and tested, the system of the RDP-II rRNA database appears to best fit current 16S rRNA sequence data for *Cyanobacteria*. The thirteen families reflect the tree topology better, and suffer less from polyphyly. Nevertheless, the CTU-based taxonomic hierarchy suggested here, further improves upon the current RDP-II system. With less higher rank level groups and more genera, our taxonomy better fits the current sequence data. Certainly, this taxonomy overlooks aspects of the polyphasic approach which also takes into account morphological and biochemical properties, or molecular markers other than 16S rRNA. Still, it is consistent, follows taxonomic group delineation rules that are not entirely subjective, and is easily extensible with new cultivated sequence data. A possible option to carry this taxonomy to real life could be to combine the species names with the numeric rank names.

### D. Applicability and limitations

Although we mainly focused on microbial 16S datasets, SATIVA can be potentially used for any clade in the tree of life and any set of genes. The only prerequisite is a reliable multiple sequence alignment of all taxa. Depending on the clade, this might become a serious obstacle if no universal marker gene is available or when aligning homologues from evolutionary distant organisms becomes challenging. Furthermore, for markers which exhibit extremely high evolution rates (like the ITS region commonly used as a fungal barcode), even genus-level alignment might be problematic (41). Conversely, slowly-evolving marker genes might have identical sequences shared by multiple closely related species. Obviously, such species would be indistinguishable for SATIVA (as for any other sequence-based approach), and thus should either be excluded from analyses or merged into a single representative pseudo-taxon. If two or more species share a certain percentage of identical sequences (default: 60%), SATIVA will issue a warning and automatically merge them into a pseudo-taxon.

A further complication is the presence of synonyms, that is, different names which refer to the same taxon (e.g., anamorphic and teleomorphic names of fungal species). To deal with this problem, SATIVA allows users to specify a list of synonymous taxon names in a separate text file. Subsequently, SATIVA will internally treat all synonyms as one single taxon.

Please note, that the presence of chimeric and/or poor-quality sequences in the alignment might seriously affect SATIVA results. Therefore, we recommend using state-of-the-art chimera detection tool (e.g., UCHIME (42)), as well as filtering/trimming methods based on sequence quality scores (like those implemented in QIIME (31)) before running SATIVA.

Another practical limitation is the running time, which grows rapidly with increasing tree size (i.e., number of sequences). In our tests, the *Cyanobacteria* phylum (≈ 1000 taxa) could be analyzed in acceptable times on a typical laptop, whereas core bacterial datasets (≈ 10000 taxa) took between 1.5 and 2 days on a multi-core node (see Section Memory consumption and running time for details). Currently, we are investigating ways to improve run-times and scalability for larger datasets, such as full 16S rRNA databases or all available GenBank sequences for a certain taxonomic rank.

For single-gene 16S rRNA alignments, the multifurcation resolution step represents a scalability bottleneck, since it cannot be parallelized efficiently with the current RAxML implementation due to the short alignment length. At the same time, the leave-one-out test is straight-forward to parallelize across taxa and thus scales well. Furthermore, our analysis of several microbial taxonomies suggests that inter-phylum mislabels are extremely rare. For higher organisms, we expect this effect to be even more pronounced, with most conflicts occurring at the lower taxonomic ranks. This also means that each phylum (or even class) can be analyzed independently, thus greatly reducing the computational burden.

Finally, we want to emphasize that SATIVA mislabel identification and re-annotation suggestions should be regarded as putative. Additional evidence including both morphological and molecular data (e.g., other marker genes) should be evaluated before taking a final decision to re-annotate. Still, our tool can yield substantial savings in man-hours by short-listing putative mislabels. Therefore, visual tree inspection, an error-prone and labor-intensive process, which can take up to several days even for an experienced taxonomist, becomes obsolete.

### E. Future directions

Apart from the aforementioned performance improvements, we plan to evaluate SATIVA in distinct settings. First, we intend to analyze marker gene databases for microbial organisms such as Fungi (19) and unicellular eukaryotes (18). Second, we consider assessing the consistency between taxonomies and phylogenies built from multiple marker genes or even whole genomes. At the same time, we believe that sequence database maintainers could benefit from using SATIVA (or similar approaches) to validate their respective classifications. In particular, the results we obtained in this study will be used to improve taxonomic annotations in the upcoming versions of the SILVA database.

Finally, we recognize that rather than correcting the errors post hoc, it would be much more efficient to prevent suboptimal taxonomic annotations from being deposited in public databases. One way to deal with this problem would be to encourage submitters to follow the best practices in sequence quality control (43). A more reliable solution could involve an automatic pre-submission plausibility check. This test can rely on fast to compute parsimony-based placements as implemented in EPA, for instance.

## V. Supplementary data

Supplementary Data are available at NAR Online.

## VII. ACKNOWLEDGMENTS

We thank Tomas Flouri, Andre Aberer and Fernando Izquierdo-Carrasco for their early work (not included in this paper) on this project. Furthermore, we would like to thank Markus Göker and Guido Grimm for their helpful comments.

The authors gratefully acknowledge the Gauss Centre for Supercomputing e.V. (www.gauss-centre.eu) for funding this project by providing computing time on the GCS Supercomputer SuperMUC at Leibniz Supercomputing Centre (LRZ, www.lrz.de).

